# GNE: A deep learning framework for gene network inference by aggregating biological information

**DOI:** 10.1101/300996

**Authors:** K C Kishan, Rui Li, Feng Cui, Qi Yu, Anne R. Haake

## Abstract

The topological landscape of gene interaction networks provides a rich source of information for inferring functional patterns of genes or proteins. However, it is still a challenging task to aggregate heterogeneous biological information such as gene expression and gene interactions to achieve more accurate inference for prediction and discovery of new gene interactions. In particular, how to generate a unified vector representation to integrate diverse input data is a key challenge addressed here. We propose a scalable and robust deep learning framework to learn embedded representations to unify known gene interactions and gene expression for gene interaction predictions. These low-dimensional embeddings derive deeper insights into the structure of rapidly accumulating and diverse gene interaction networks and greatly simplify downstream modeling. We compare the predictive power of our deep embeddings to the strong baselines. The results suggest that our deep embeddings achieve significantly more accurate predictions. Moreover, a set of novel gene interaction predictions are validated by up-to-date literature-based database entries. GNE is freely available under the GNU General Public License and can be downloaded from Github (https://github.com/kckishan/GNE)

## 1 Background

A comprehensive study of gene interactions (GIs) provides means to identify the functional relationship between genes and their corresponding products, as well as insights into underlying biological phenomena that are critical to understanding phenotypes in health and disease conditions^1–3^. Since advancements in measurement technologies have led to numerous high-throughput datasets, there is a great value in developing efficient computational methods capable of automatically extracting and aggregating meaningful information from heterogeneous datasets to infer gene interactions.

Although a wide variety of machine learning models have been developed to analyze high-throughput datasets for GI prediction^4^, there are still some major challenges, such as efficient analysis of large heterogeneous datasets, integration of biological information, and effective feature engineering. To address these challenges, we propose a novel deep learning framework to integrate diverse biological information for GI network inference.

Our proposed method frames GI network inference as a problem of network embedding. In particular, we represent gene interactions as a network of genes and their interactions and create a deep learning framework to automatically learn an informative representation which integrates both the topological property and the gene expression property. A key insight behind our gene network embedding method is the "guilt by association" assumption^5^, that is, genes that are co-localized or have similar topological roles in the interaction network are likely to be functionally correlated. This insight not only allows us to discover similar genes and proteins but also to infer the properties of unknown ones. Our network embedding generates a lower-dimensional vector representation of the gene topological characteristics. The relationships between genes including higher-order topological properties are captured by the distances between genes in the embedding space. The new low-dimensional representation of a GI network can be used for various downstream tasks, such as gene function prediction, gene interaction prediction, and gene ontology reconstruction^6^.

Furthermore, since the network embedding method can only preserve the topological properties of a GI network, and fails to generalize for genes with no interaction information, our scalable deep learning method also integrates heterogeneous gene information, such as expression data from high throughput technologies, into the GI network inference. Our method projects genes with similar attributes closer to each other in the embedding space, even if they may not have similar topological properties. The results show that by integrating additional gene information in the network embedding process, the prediction performance is improved significantly.

GI prediction is a long-standing problem. The proposed machine learning methods include statistical correlation, mutual information^7^, dimensionality reduction^8^, and network-based methods (e.g. common neighborhood, network embedding)^4,9^. Among these methods, some methods such as statistical correlation and mutual information consider only gene expression whereas other methods use only topological properties to predict GIs.

Network-based methods have been proposed to leverage topological properties of GI networks^10^. Neighborhood-based methods quantify the proximity between genes, based on common neighbors in GI network^11^. The proximity scores assigned to a pair of genes rely on the number of neighbors that the pair has in common. Adjacency matrix, representing the interaction network, or proximity matrix, obtained from neighborhood-based methods, are processed with network embedding methods to learn embeddings that preserve the structural properties of the network. Structure-preserving network embedding methods such as Isomap^12^ are proposed as a dimensionality reduction technique. Since the goal of these methods is solely for graph reconstruction, the embedding space may not be suitable for GI network inference. Besides, these methods construct the graphs from the data features where proximity between genes is well defined in the original feature space^9^. On the other hand, in GI networks, gene proximities are not explicitly defined, and they depend on the specific analytic tasks and application scenarios.

Our deep learning method allows incorporating gene expression data with GI network topological structure information to preserve both topological and attribute proximity in the low-dimensional representation for GI predictions. Moreover, the scalable architecture enables us to incorporate additional attributes. Topological properties of GI network and expression profiles are transformed into two separate embeddings: ID embedding (which preserves the topological structure proximity) and attribute embedding (which preserves the attribute proximity) respectively. With a multilayer neural network, we then aggregate the complex statistical relationships between topology and attribute information to improve GI predictions.

In summary, our contributions are as follows:

- We propose a novel deep learning framework to learn lower dimensional representations while preserving topological and attribute proximity of GI networks.
- We evaluate the prediction performance on the datasets of two organisms based on the embedded representation and achieve significantly better predictions than the strong baselines.
- Our method can predict new gene interactions which are validated on an up-to-date GI database.

### Methods

#### Preliminaries

We formally define the problem of gene network inference as a network embedding problem using the concepts of topological and attribute proximity as demonstrated in Figure 1.

**Figure 1.**
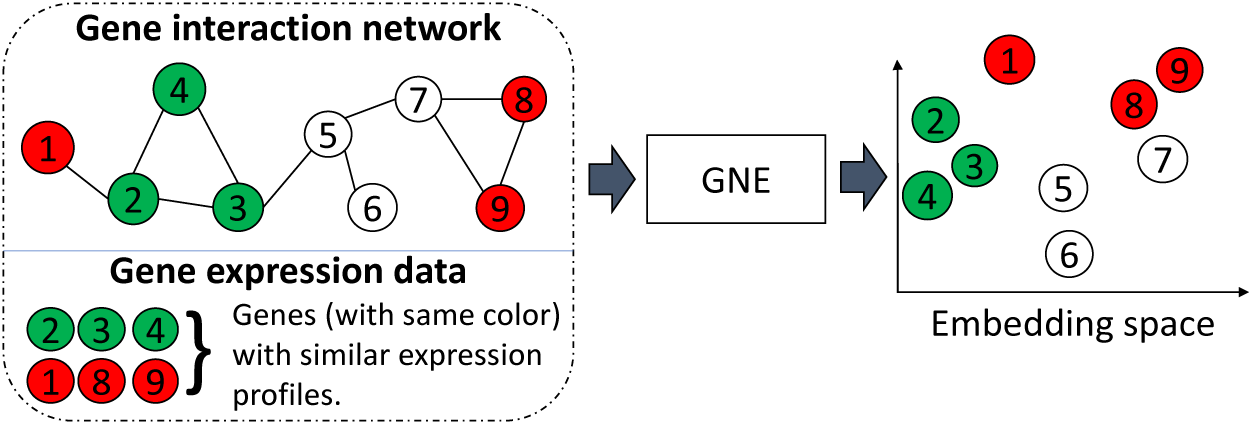
An illustration of gene network embedding (GNE). GNE integrates gene interaction network and gene expression data to learn a lower-dimensional representation.The nodes represent genes, and the genes with the same color have similar expression profiles. GNE groups genes with similar network topology, which are connected or have a similar neighborhood in the graph, and attribute similarity (similar expression profiles) in the embedded space.

Definition 1: (**Gene network**) Gene network can be represented as a network structure, which represents the interactions between genes within an organism. The interaction between genes corresponds to either a physical interaction through their gene products, e.g., proteins, or one of the genes alters or affects the activity of other gene of interest. We denote gene network as *G* = (*V,E,A*) where *V* = {*v*_*i*_} denotes genes or proteins, *E* = {*e*_*ij*_} denotes edges that correspond to interactions between genes *v*_*i*_ and *v*_*j*_, and *A* = {*A*_*i*_} represents the attributes of gene *v*_*i*_. Edge *e*_*ij*_ is associated with a weight *w*_*ij*_ ≥ 0 indicating the strength of the connection between gene *v*_*i*_ and *v*_*j*_. If gene *v*_*i*_ and *v*_*j*_ is not linked by an edge, *w*_*ij*_ = 0. We name interactions with *w*_*ij*_ > 0 as positive interactions and *w*_*ij*_ = 0 as negative interactions. In this paper, we consider weights *w*_*ij*_ to be binary, indicating whether genes *v*_*i*_ and *v*_*j*_ interact or not.

Genes directly connected with a gene *v*_*i*_ in gene network denote the local network structure of gene *v*_*i*_. We define local network structures as the first-order proximity of a gene.

Definition 2: (**First-order proximity**) The first-order proximity in a gene network is the pairwise interactions between genes. Weight *w*_*ij*_ indicates the first-order proximity between gene *v*_*i*_ and *v*_*j*_. If there is no interaction between gene *v*_*i*_ and *v*_*j*_, their first-order proximity *w*_*ij*_ is 0.

Genes are likely to be involved in the same cellular functions if they are connected in the gene network. On the other hand, even if two genes are not connected, they may be still related in some cellular functions. This indicates the need for an additional notion of proximity to preserve the network structure. Studies suggest that genes that share a similar neighborhood are also likely to be related^6^. Thus, we introduce second-order proximity that characterizes the global network structure of the genes.

Definition 3: (**Second-order proximity**) Second order proximity denotes the similarity between the neighborhood of genes. Let *N*_*i*_ = {*s*_*i*_,_1_,…, *s*_*i,i*−1_, *s*_*i,i*+1_,…, *s*_*i,M*−1_} denotes the first-order proximity of gene *v*_*i*_, where *s*_*i,j*_ is *w*_*ij*_ if there is direct connection between gene *v*_*i*_ and gene *v*_*j*_, otherwise 0. Then, the second order proximity is the similarity between *N*_*i*_ and *N*_*j*_. If there is no path to reach gene *v*_*i*_ from gene *v*_*j*_, the second proximity between these genes is 0.

Integrating first-order and second-order proximities simultaneously can help to preserve the topological properties of the gene network. To generate a more comprehensive representation of the genes, it is crucial to integrate gene expression data as gene attributes with their topological properties. Besides preserving topological properties, gene expression provides additional information to predict the network structure.

Definition 4: (**Attribute proximity**) Attribute proximity denotes the similarity between the expression of genes.

We thus investigate both topological and attribute proximity for gene network embedding, which is defined as follows:

Definition 5: (**Gene network embedding**) Given a gene network denoted as *G* = (*V,E,A*), gene network embedding aims to learn a function *f* that maps gene network structure and their attribute information to a *d*-dimensional space where a gene is represented by a vector *y*_*i*_ ∈ ℝ^*d*^ where *d* ≪ *M*. The low dimensional vectors *y*_*i*_ and *y*_*j*_ for genes *v*_*i*_ and *v*_*j*_ preserve their relationships in terms of the network topological structure and attribute proximity.

#### Gene Network Embedding (GNE) model

Our deep learning framework as shown in Figure 2 jointly utilizes gene network structure and gene expression data to learn a unified representation for the genes. Embedding of a gene network projects genes into a lower dimensional space, known as the embedding space, in which each gene is represented by a vector. The embeddings preserve both the gene network structure and statistical relationships of gene expression. We list the variables to specify our framework in Table 1.

**Figure 2.**
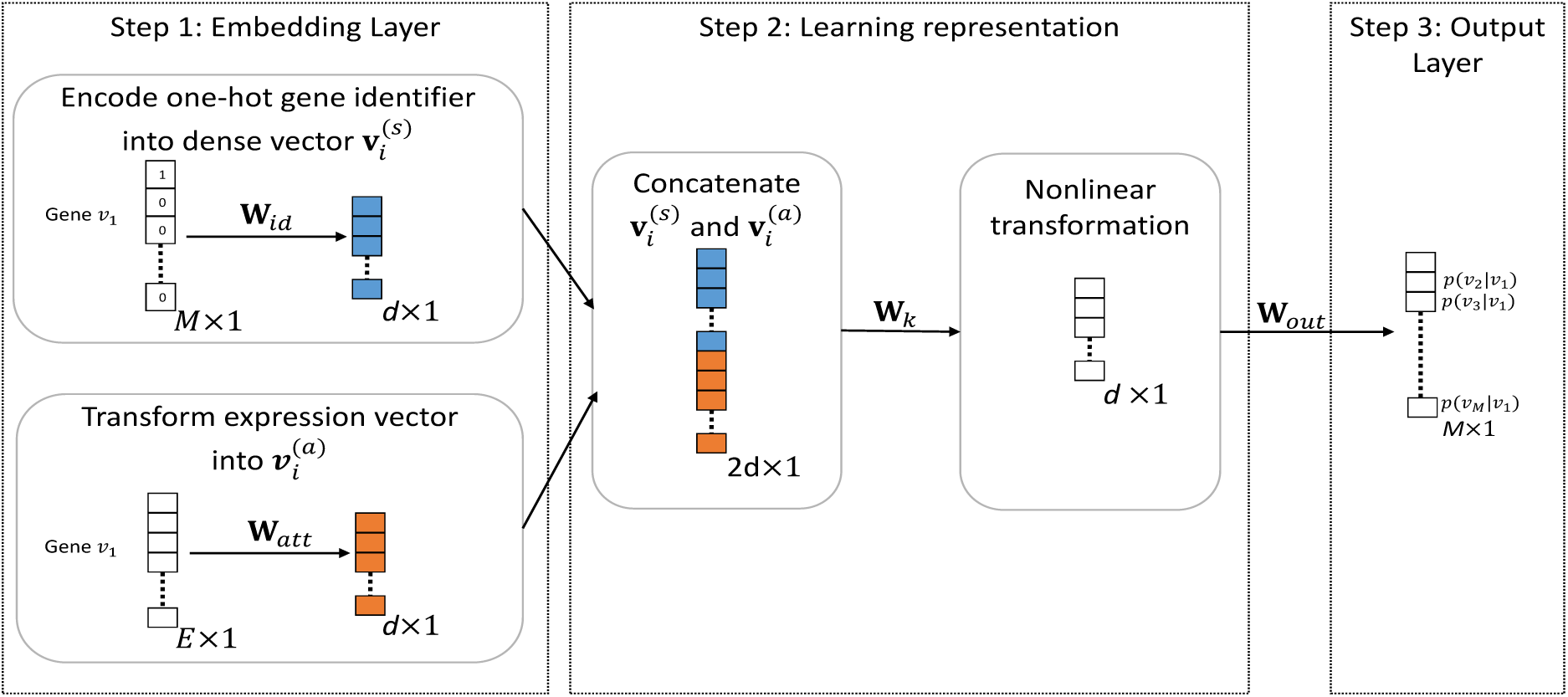
Overview of Gene Network Embedding (GNE) Framework for gene interaction prediction. On the left,one-hot encoded representation of gene is encoded to dense vector 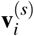 of dimension *d* × 1 which captures topological properties and expression vector of gene is transformed to 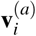 of dimension *d* × 1 which aggregates the attribute information (Step 1). Next, concatenation of two embedded vectors (creates vector with dimension 2*d* × 1) allows to combine strength of both network structure and attribute modeling. Then, nonlinear transformation of concatenated vector enables GNE to capture complex statistical relationships between network structure and attribute information and learn better representations (Step 2). Finally, these learned representation of dimension *d* × 1 is transformed into a probability vector of length *M* × 1 in output layer, which contains the predictive probability of gene *v*_*i*_ to all the genes in the network. Conditional probability *p*(*v*_*j*_|*v*_*i*_) on output layer indicates the likelihood that gene *v*_*j*_ is connected with gene *v*_*i*_ (Step 3).

**Table 1.** Terms and Notations

| Symbol | Definitions |
| --- | --- |
| $M$ | Total number of genes in gene network |
| $E$ | Number of expression values for each gene |
| $N_i$ | Set of the neighbor genes of gene $v_i$ |
| $\mathbf{v}_i^{(s)}$ | Topological representation of gene $v_i$ |
| $\mathbf{v}_i^{(a)}$ | Attribute representation of gene $v_i$ |
| $\tilde{\mathbf{v}}_i$ | Neighborhood representation of gene $v_i$ |
| $\mathbf{v}_i$ | Concatenated representation of topological properties and expression data |
| $k$ | Number of hidden layers to transform concatenated representation into embedding space |
| $\mathbf{h}^{(k)}$ | Output of $k^{th}$ hidden layer |
| $\mathbf{W}_k$ | Weight matrix for $k^{th}$ hidden layer |
| $\mathbf{W}_{id}$ | Weight matrix for topological structure embedding |
| $\mathbf{W}_{att}$ | Weight matrix for attribute embedding |
| $\mathbf{W}_{out}$ | Weight matrix for output layer |

##### Gene Network Structure Modeling

GNE framework preserves first-order and second-order proximity of genes in the gene network. The key idea of network structure modeling is to estimate the pairwise proximity of genes in terms of the network structure. If two genes are connected or share similar neighborhood genes, they tend to be related and should be placed closer to each other in the embedding space. Inspired by the Skip-gram model^13^, we use one hot encoded representation to represent topological information of a gene. Each gene *v*_*i*_ in the network is represented as an *M*-dimensional vector where only the *i*^*th*^ component of the vector is 1.

To model topological similarity, we define the conditional probability of gene *v*_*j*_ on gene *v*_*i*_ using a softmax function as:

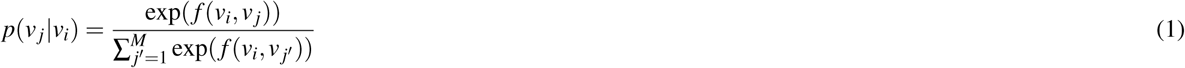

which measures the likelihood of gene *v*_*i*_ being connected with *v*_*j*_. Let function *f* represents the mapping of two genes *v*_*i*_ and *v*_*j*_ to their estimated proximity score. Let *p*(*N|v*) be the likelihood of observing a neighborhood *N* for a gene *v*. By assuming conditional independence, we can factorize the likelihood so that the likelihood of observing a neighborhood gene is independent of observing any other neighborhood gene, given a gene *v*_*i*_:

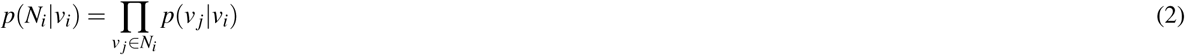

where *N*_*i*_ represents the neighborhood genes of the gene *v*_*i*_. Global structure proximity for a gene *v*_*i*_ can be preserved by maximizing the conditional probability over all genes in the neighborhood. Hence, we can define the likelihood function that preserve global structure proximity as:

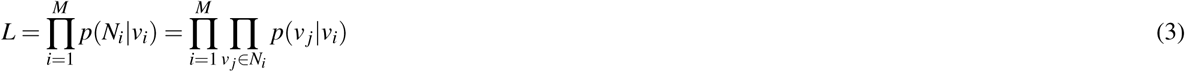

Let 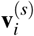 denotes the dense vector generated from one-hot gene ID vector, which represents topological information of that gene. GNE follows direct encoding methods^13,14^ to map genes to vector embeddings, simply known as embedding lookup:

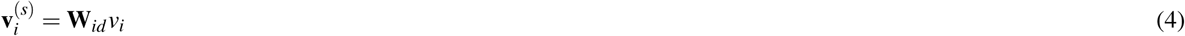

where **W**_*id*_ ∈ ℝ^*d×M*^ is a matrix containing the embedding vectors for all genes and 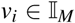 indicating the column of **W**_*id*_ corresponding to gene *v*_*i*_.

##### Gene Expression Modeling

GNE encodes the expression data from microarray experiments to the dense representation using a non-linear transformation. The amount of mRNA produced during transcription measured over a number of experiments helps to identify similarly expressed genes. Since expression data have inherent noise^15^, transforming expression data using a non-linear transformation can be helpful to uncover the underlying representation. Let *x*_*i*_ be the vector of expression values of gene *v*_*i*_ measured over *E* experiments. Using non-linear transformation, we can capture the non-linearities of expression data of gene *v*_*i*_ as:

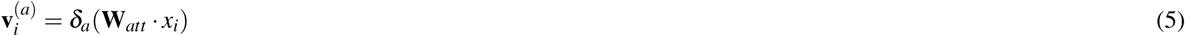

where 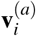 represents the lower dimensional attribute representation vector for gene *v*_*i*_. **W**_*att*_, and *δ*_*a*_ represents the weight matrix, and activation function of attribute transformation layer respectively.

We use the deep model to approximate the attribute proximity by capturing complex statistical relationships between attributes and introducing non-linearities, similar to structural embedding.

##### GNE Integration

GNE models the integration of network structure and attribute information to learn more comprehensive embeddings for gene networks. GNE takes two inputs: one for topological information of a gene as one hot gene ID vector and another for its expression as an attribute vector. Each input is encoded to its respective embeddings. One hot representation for a gene *v*_*i*_ is projected to the dense vector 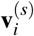 which captures the topological properties. Non-linear transformation of attribute vector generates compact representation vector 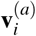. Previous work^16^ combines heterogeneous information using the late fusion approach. However, the late fusion approach is the approach of learning separate models for heterogeneous information and integrating the representations learned from separate models. On the other hand, the early fusion combines heterogeneous information and train the model on combined representations^17^. We thus propose to use the early fusion approach to combine them by concatenating. As a result, learning from topological and attribute information can complement each other, allowing the model to learn their complex statistical relationships as well. Embeddings from topological and attribute information are concatenated into a vector as:

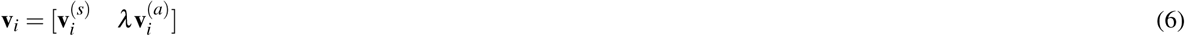

where *λ* is the importance of gene expression information relative to topological information.

The concatenated vectors are fed into a multilayer perceptron with *k* hidden layers. The hidden representations from each hidden layer in GNE are denoted as 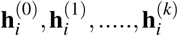, which can be defined as :

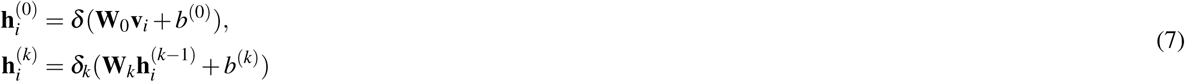

where *δ*_*k*_ represents the activation function of layer *k*. 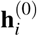 represents initial representation and 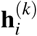 represents final representation of the input gene *v*_*i*_. Transformation of input data using multiple non-linear layers has shown to improve the representation of input data^18^. Moreover, stacking multiple layers of non-linear transformations can help to learn high-order statistical relationships between topological properties and attributes.

At last, final representation 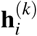 of a gene *v*_*i*_ from the last hidden layer is transformed to probability vector, which contains the conditional probability of all other genes to *v*_*i*_:

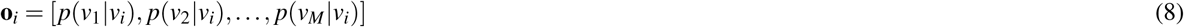

where *p*(*v*_*j*_|*v*_*i*_) represents the probability of gene *v*_*i*_ being related to gene *v*_*j*_ and **o**_*i*_ represents the output probability vector with the conditional probability of gene *v*_*i*_ being connected to all other genes.

Weight matrix **W**_*out*_ between the last hidden layer and the output layer corresponds to the abstractive representation of neighborhood of genes. A *j*^*th*^ row from **W**_*out*_ refers to the compact representation of neighborhood of gene *v*_*j*_, which can be denoted as 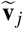. The proximity score between gene *v*_*i*_ and *v*_*j*_ can be defined as:

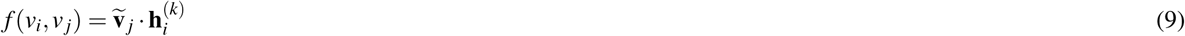

which can be replaced into Eq. 1 to calculate the conditional probability:

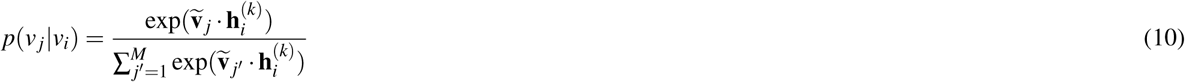

Our model learns two latent representations 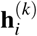 and 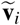 for a gene *v*_*i*_ where 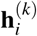 is the representation of gene as a node and 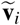 is the representation of the gene *v*_*i*_ as a neighbor. Neighborhood representation 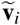 can be combined with node representation 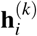 by addition^19,20^ to get final representation for a gene as:

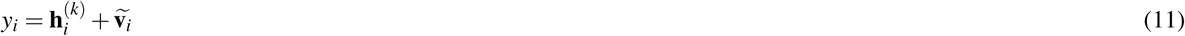

which returns us better performance results.

For an edge connecting gene *v*_*i*_ and *v*_*j*_, we create feature vector by combining embeddings of those genes using Hadamard product. Empirical evaluation shows features created with Hadamard product gives better performance over concatenation^14^. Then, we train a logistic classifier on these features to classify whether genes *v*_*i*_ and *v*_*j*_ interact or not.

##### Parameter Optimization

To optimize GNE, the goal is to maximize objective function mentioned in Eq. 10 as a function of all parameters. Let Θ be the parameters of GNE which includes {**W**_*id*_,**W**_*att*_,**W**_*out*_,Θ_*h*_} and Θ_*h*_ represents weight matrices **W**_*k*_ of hidden layers. We train our model to maximize the objective function with respect to all parameters Θ :

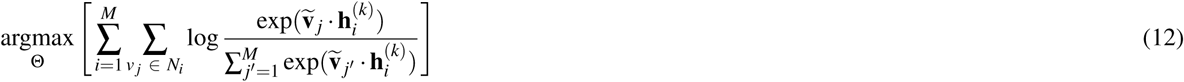

Maximizing this objective function with respect to Θ is computationally expensive, which requires the calculation of partition function 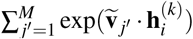 for each gene. To calculate a single probability, we need to aggregate all genes in the network. To address this problem, we adopt the approach of negative sampling^13^ which samples the negative interactions, interactions with no evidence of their existence, according to some noise distribution for each edge *e*_*ij*_. This approach allows us to sample a small subset of genes from the network as negative samples for a gene, considering that the genes on selected subset don’t fall in the neighborhood *N*_*i*_ of the gene. Above objective function enhances the similarity of a gene *v*_*i*_with its neighborhood genes *v*_*j*_ ∈ *N*_*i*_ and weakens the similarity with genes not in its neighborhood genes *v*_*j*_ ∉ *N*_*i*_. It is inappropriate to assume that the two genes in the network are not related if they are not connected. It may be the case that there is not enough experimental evidence to support that they are related yet. Thus, forcing the dissimilarity of a gene with all other genes, not in its neighborhood *N*_*i*_ seems to be inappropriate.

We adopt Adaptive Moment Estimation (Adam) optimization^21^, which is an extension to stochastic gradient descent, for optimizing Eq. 12. Adam computes the adaptive learning rate for each parameter by performing smaller updates for the frequent parameters and larger updates for the infrequent parameters. The Adam method provides the ability of AdaGrad^22^ to deal with sparse gradients and also the ability of RMSProp^23^ to deal with non-stationary objectives. In each step, Adam algorithm samples mini-batch of interactions and then updates GNE’s parameters. To address the issue of overfitting, regularization like dropout^24^ and batch normalization^25^ is added to hidden layers. Proper optimization of GNE gives the final representation for each gene.

#### Experimental setup

We evaluate our model using two real organism datasets. We take gene interaction network data from the BioGRID database^26^ and gene expression data from DREAM5 challenge^7^. We use two interaction datasets from BioGRID database (2017 released version 3.4.153 and 2018 released version 3.4.158) to evaluate the predictive performance of our model. Self-interactions and redundant interactions are removed from interaction datasets. The statistics of the datasets are shown in Table 2.

**Table 2.** Statistics of the interaction datasets from BioGRID and the gene expression data from DREAM5 challenge.

| Datasets | #(Genes) | #(Interactions) |  | Expression data |
| --- | --- | --- | --- | --- |
|  |  | 2017 version | 2018 version | #(Experiments) |
| Yeast | 5,950 | 544,652 | 557,487 | 536 |
| E. coli | 4,511 | 148,340 | 159,523 | 805 |

We evaluate the learned embeddings to infer gene network structure. We randomly hold out a fraction of interactions as the validation set for hyper-parameter tuning. Then, we divide the remaining interactions randomly into training and testing dataset with the equal number of interactions. Since the validation set and the test set contains only positive interactions, we randomly sample an equal number of gene pairs from the network, considering the missing edge between the gene pairs represents the absence of interactions. Given the gene network G with a fraction of missing interactions, the task is to predict these missing interactions.

We compare the GNE model with five competing methods. Correlation directly predicts the interactions between genes based on the correlation of expression profiles. Then, the following three baselines (Isomap, LINE, and node2vec) are network embedding methods. Specifically, node2vec is the strong baseline for structural network embedding. We evaluate the performance of GNE against the following methods:

- **Correlation^27^** It computes Pearson’s correlation coefficient between all genes and the interactions are ranked via correlation scores, i.e., highly correlated gene pairs receive higher confidence.
- **Isomap^10^** It computes all-pairs shortest-path distances to create a distance matrix and performs singular-value decomposition of that matrix to learn a lower-dimensional representation. Genes separated by the distance less than threshold *ε* in embedding space are considered to have the connection with each other and the reliability index, a likelihood indicating the interaction between two genes, is computed using FSWeight^28^.
- **LINE^16^** Two separate embeddings are learned by preserving first-order and second-order proximity of the network structure respectively. Then, these embeddings are concatenated to get final representations for each node.
- **node2vec^14^** It learns the embeddings of the node by applying Skip-gram model to node sequences generated by a biased random walk. We tuned two hyper-parameters p and q that control the random walk.

Note that the competing methods such as Isomap, LINE, and node2vec are designed to capture only the topological properties of the network. For the fair comparison with GNE that additionally integrates expression data, we concatenate attribute feature vector with learned gene representation to extend baselines by including the gene expression. We name these variants as Isomap+, LINE+, and node2vec+.

We have implemented GNE with TensorFlow framework^29^. The parameter settings for GNE are determined by its performance on the validation set. We randomly initialize GNE’s parameters, optimizing with mini-batch Adam. We test the batch size of [8, 16, 32, 64, 128, 256] and learning rate of [0.1, 0.01, 0.005, 0.002, 0.001, 0.0001]. We test the number of negative samples to be [2, 5, 10, 15, 20] as suggested by^13^. We test the embedding dimension *d* of [32, 64, 128, 256] for all methods. Also, we evaluate model’s performance with respect to different values of *λ* [0, 0.2, 0.4, 0.6, 0.8, 1], which is discussed in more detail later. The parameters are selected based on empirical evaluation and Table 3 summarizes the optimal parameters tuned on validation data sets.

**Table 3.** Optimal parameter settings for GNE model

| Dataset | Learning rate | Batch size | Embedding dimension (d) | Epoch | Negative samples |
| --- | --- | --- | --- | --- | --- |
| Yeast | 0.005 | 256 | 128 | 20 | 10 |
| E. coli | 0.002 | 128 | 128 | 20 | 10 |

To capture the non-linearity of gene expression data, we choose Exponential Linear Unit (ELU)^30^ activation function, which corresponds to *δ*_*a*_ in Eq. 5. Also, ELU activation avoids vanishing gradient problem and provides improved learning characteristics in comparison to other methods. We use a single hidden layer (*k* = 1) with hyperbolic tangent activation (Tanh) to model complex statistical relationships between topological properties and attributes of the gene. The choice of ELU for attribute transformation and Tanh for hidden layer shows better performance upon empirical evaluation.

We use the area under the ROC curve (AUROC) and area under the precision-recall curve (AUPR)^31^ to evaluate the rankings generated by the model for interactions in the test set. These metrics are widely used in evaluating the ranked list of predictions in gene interaction^4^.

### Results and Discussion

We evaluate the ability of our GNE model to predict gene interaction of two real organisms. We present empirical results of our proposed method against other methods.

#### Analysis of gene embeddings

We visualize the embedding vectors of genes learned by GNE. We take the learned embeddings, which specifically model the interactions by preserving topological and attribute similarity. We embed these embeddings into a 2D space using t-SNE package^32^ and visualize them (Figure 3). For comparison, we also visualize the embeddings learned by structure-preserving deep learning methods, such as LINE, and node2vec.

**Figure 3.**
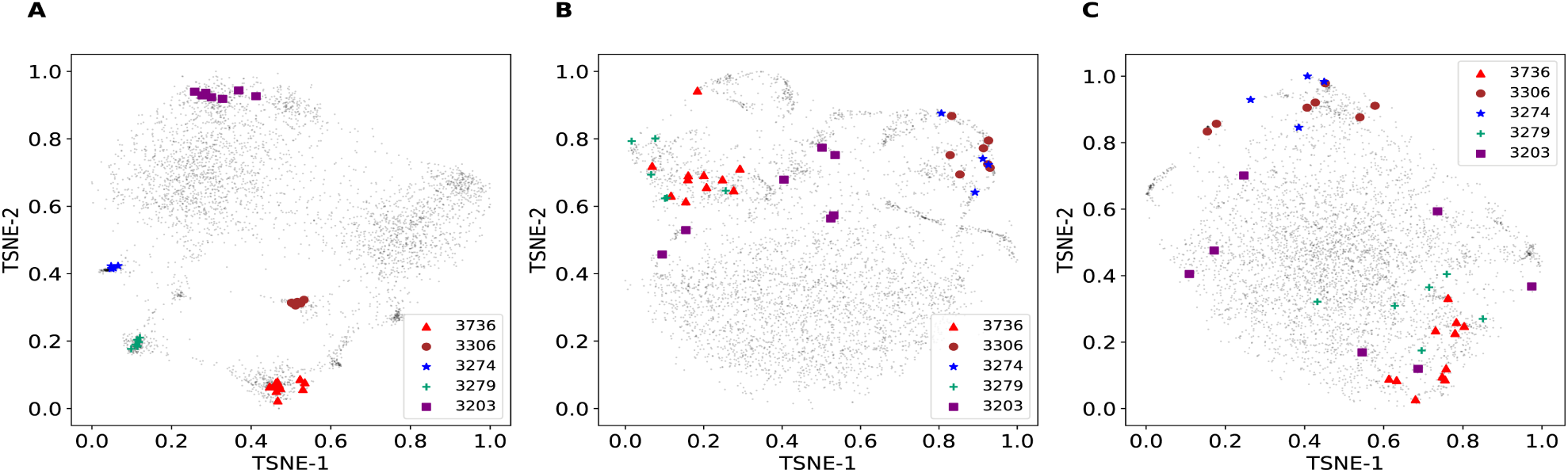
Visualization of learned embeddings for genes on E. coli. Genes are mapped to the 2D space using the t-SNE package^32^ with learned gene representations (*y*_*i*_,*i* = 1,2*,…,M*) from different methods: (A) GNE, (B) LINE, and (C) node2vec as input. Operons 3203, 3274, 3279, 3306, and 3736 of E. coli are visualized and show clustering patterns. Best viewed on screen.

In E. coli, a substantial fraction of functionally related genes are organized into operons, which are the group of genes that interact with each other and are co-regulated^33^. Since this concept fits well with the topological and attribute proximity implemented in GNE, we expect GNE to place genes within an operon close to each other in the embedding space. To evaluate this, we collect information about operons of E. coli from the DOOR database and visualize the embeddings of genes within these operons (Figure 3).

Figure 3 reveals the clustering structure that corresponds to the operons on E. coli. For example, operon with operon id 3306 consists of seven genes: rsxA, rsxB, rsx, rsxD, rsxG, rsxE, and nth that are involved in electron transport. GNE infers similar representations for these genes, resulting in localized projection in the 2D space. Similarly, other operons also show similar patterns (Figure 3).

To test if the pattern in Figure 3 holds across all operons, we compute the average Euclidean distance between each gene’s vector representation and vector representations of other genes within the same operon. Genes within the same operon have significantly similar vector representation *y*_*i*_ than expected by chance (p-value = 1.75*e* − 127, 2-sample KS test).

Thus, the analysis here indicates that GNE can learn similar representations for genes with similar topological properties and expression.

#### Gene Interaction Prediction

We randomly remove 50% of interactions from the network and compare various methods to evaluate their predictions for 50% missing interactions. Table 4 shows the performance of GNE and other methods on gene interaction prediction across different datasets. As our method significantly outperforms other competing methods, it indicates the informativeness of gene expression in predicting missing interactions. Also, our model is capable of integrating attributes with topological properties to learn better representations.

**Table 4.** Area under ROC curve (AUROC) and Area under PR curve (AUPR) for gene Interaction Prediction. + indicates the concatenation of expression data with learned embeddings to create final representation. **\*** denotes that GNE significantly outperforms node2vec at 0.01 level paired t-test. Note that method that achieves the best performance is bold faced.

| Methods | Yeast |  | E. coli |  |
| --- | --- | --- | --- | --- |
|  | AUROC | AUPR | AUROC | AUPR |
| Correlation | 0.582 | 0.579 | 0.537 | 0.557 |
| Isomap | 0.507 | 0.588 | 0.559 | 0.672 |
| LINE | 0.726 | 0.686 | 0.897 | 0.851 |
| node2vec | 0.739 | 0.708 | 0.912 | 0.862 |
| Isomap+ | 0.653 | 0.652 | 0.644 | 0.649 |
| LINE+ | 0.745 | 0.713 | 0.899 | 0.856 |
| node2vec+ | 0.751 | 0.716 | 0.871 | 0.826 |
| GNE (Topology) | 0.787 | 0.784 | 0.930 | 0.931 |
| **GNE (our model)** | **0.825\*** | **0.821\*** | **0.940\*** | **0.939\*** |

We compare our model with a correlation-based method, that takes only expression data into account. Our model shows significant improvement of 0.243 (AUROC), 0.242 (AUPR) on yeast and 0.403 (AUROC), 0.382 (AUPR) on E. coli over correlation-based methods. This improvement suggests the significance of the topological properties of the gene network.

The network embedding method, Isomap, performs poorly in comparison to correlation-based methods on yeast because of its limitation on network inference. Deep learning based network embedding methods such as LINE, and node2vec show the significant gain over Isomap and correlation-based methods. node2vec outperforms LINE across two datasets. Moreover, GNE trained only with topological properties outperforms these structured-based deep learning methods (Table 4). However, these methods don’t consider the attributes of the gene that we suggest to contain useful information for gene interaction prediction. By adding expression data with topological properties, GNE outperforms structure-preserving deep embedding methods across both datasets.

Focusing on the results corresponding to the integration of expression data with topological properties, we find that the method of integrating the expression data plays an essential role in the performance. Performance of node2vec+ (LINE+, Isomap+) shows little improvement with the integration of expression data on yeast. However, node2vec+ (LINE+, Isomap+) has no improvement or decline in performance on E. coli. The decline in performance indicates that merely concatenating the expression vector with learned representations for the gene is insufficient to capture the rich information in expression data. The late fusion approach of combining the embedding vector corresponding to the topological properties of the gene network and the feature vector representing expression data has no significant improvement in the performance (except Isomap). In contrast, our model incorporates gene expression data with topological properties by the early fusion method and shows significant improvement over other methods.

#### Impact of network sparsity

We investigate the robustness of our model to network sparsity. We hold out 10% interactions as the test set and change the sparsity of the remaining network by randomly removing a portion of remaining interactions. Then, we train GNE to predict interactions in the test set and evaluate the change in performance to network sparsity. We evaluate two versions of our implementations: GNE with only topological properties and GNE with topological properties and expression data. The result is shown in Figure 4.

**Figure 4.**
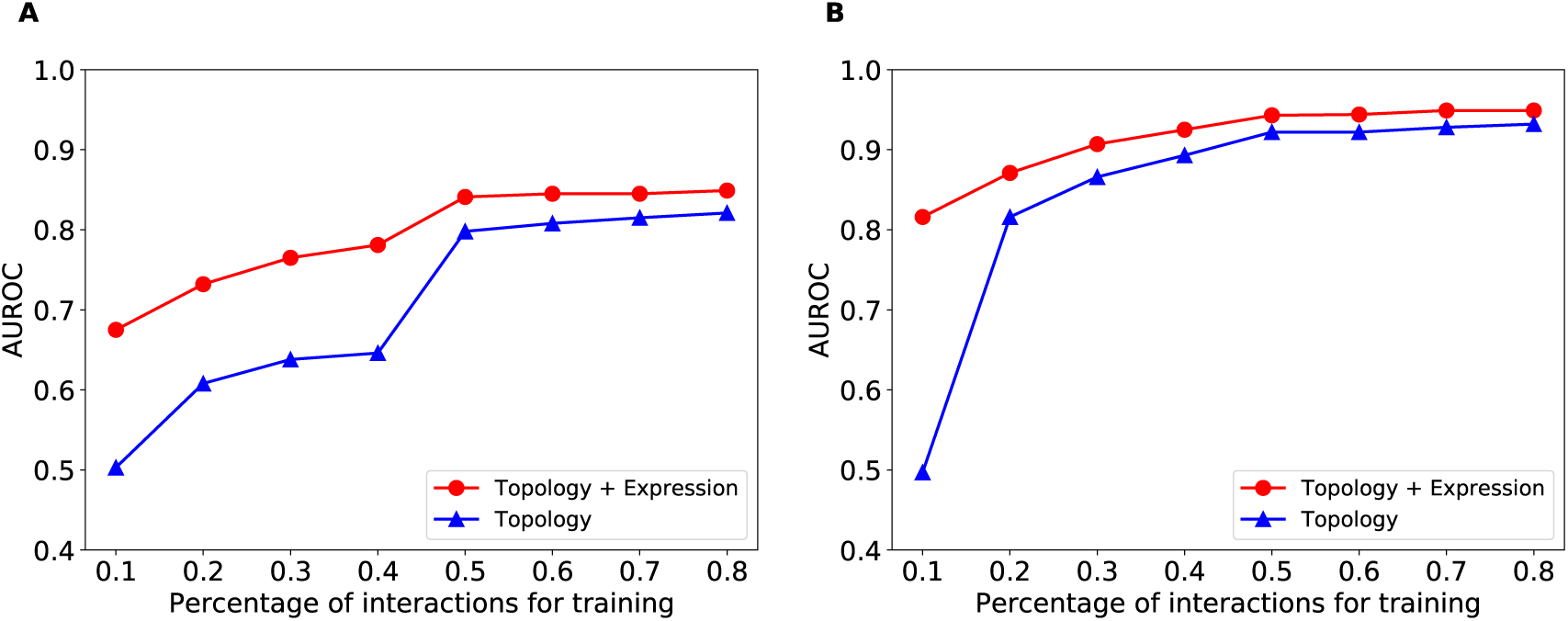
AUROC comparison of GNE’s performance with respect to network sparsity. (A) yeast (B) E. coli. Integration of expression data with topological properties of the gene network improves the performance for both datasets.

Figure 4 shows that our method’s performance improves with an increase in the number of training interactions across datasets. Also, our method’s performance improves when expression data is integrated with the topological structure. Specifically, GNE trained on 10% of total interactions and attributes of yeast shows a significant gain of 0.172 AUROC (from 0.503 to 0.675) over GNE trained only with 10% of total interactions as shown in Figure 4. Similarly, GNE improves the AUROC from 0.497 to 0.816 for E. coli with the same setup as shown in Figure 4. The integration of gene expression data results in less improvement when we train GNE on a relatively large number of interactions.

Moreover, the performance of GNE trained with 50% of total interactions and expression data is comparable to be trained with 80% of total interactions without gene expression data as shown in Figure 4. The integration of expression data with topological properties into GNE model has more improvement on E. coli than yeast when we train with 10% of total interactions for each dataset. The reason for this is likely the difference in the number of available interactions for yeast and E. coli (Table 2). This indicates the informativeness of gene expression when we have few interactions and supports the idea that the integration of expression data with topological properties improves gene interaction prediction.

#### Impact of *λ*

GNE involves the parameter *λ* that controls the importance of gene expression information relative to topological properties of gene network as shown in Eq. 6. We examine how the choice of the parameter *λ* affects our method’s performance. Figure 5 shows the comparison of our method’s performance with different values of *λ* when GNE is trained on varying percentage of total interactions.

**Figure 5.**
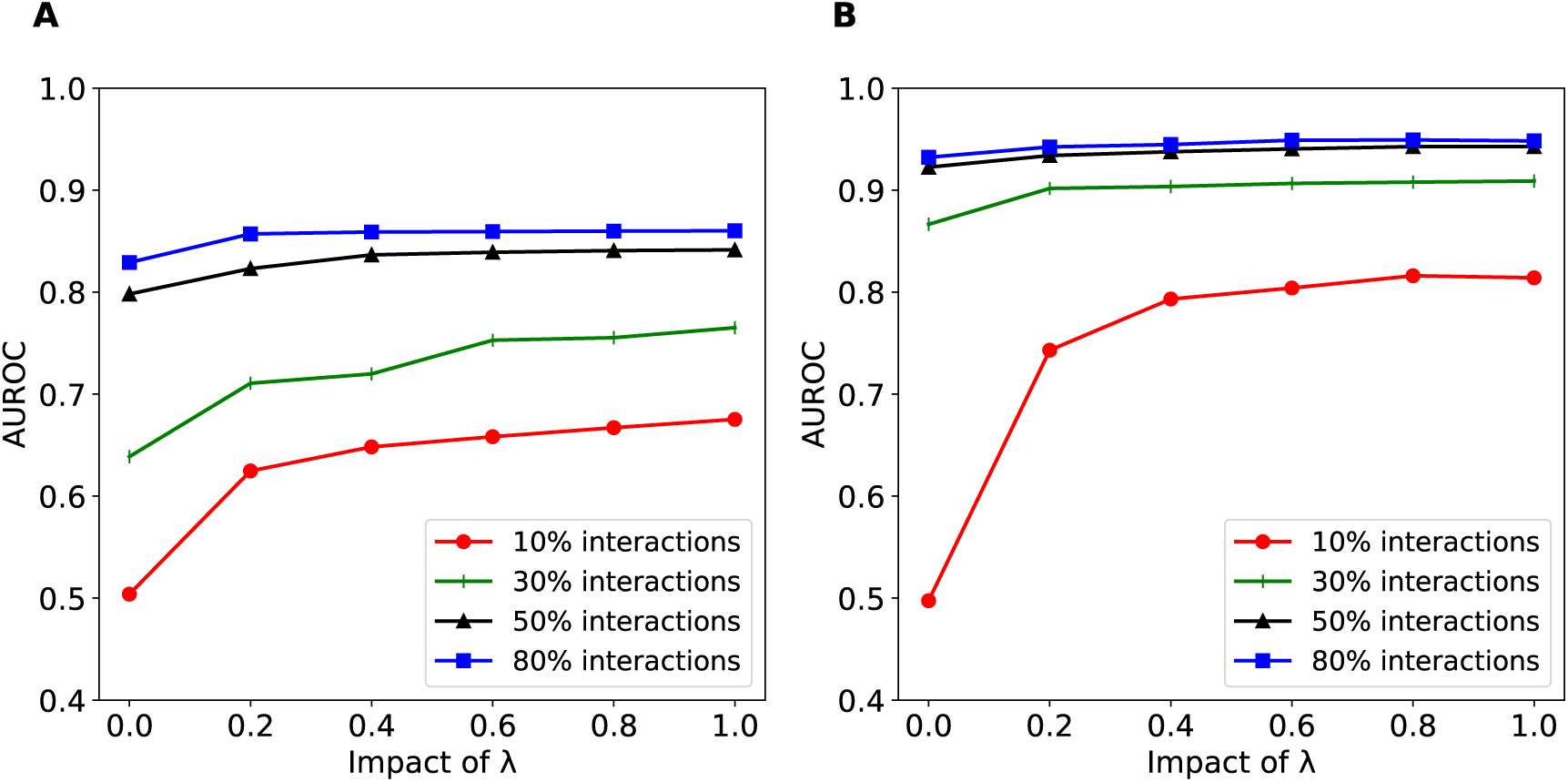
Impact of *λ* on GNE’s performance trained with different percentages of interactions. (A) yeast (B) E. coli. Different lines indicate performance of GNE trained with different percentages of interactions.

We evaluate the impact of *λ* on range [0, 0.2, 0.4, 0.6, 0.8, 1]. When *λ* becomes 0, the learned representations model only topological properties. In contrast, setting the high value for *λ* makes GNE learn only from attributes and degrades its performance. Therefore, our model performs well when *λ* is within [0, 1].

Figure 5 shows that the integration of expression data improves the performance of GNE to predict gene interactions. Impact of *λ* depends on the number of interactions used to train GNE. If GNE is trained with few interactions, integration of expression data with topological properties plays a vital role in predicting missing interactions. As the number of training interactions increases, integration of expression data has less impact but still improves the performance over only topological properties.

Figure 4 and 5 demonstrate that the expression data contributes the increase in AUROC by nearly 0.14 when interactions are less than 40% for yeast and about 0.32 when interactions are less than 10% for E. coli. More topological properties and attributes are required for yeast than E. coli. It may be related to the fact that yeast is a more complex species than E. coli. Moreover, we can speculate that more topological properties and attributes are required for higher eukaryotes like humans. In humans, GNE that integrates topological properties with attributes may be more successful than the methods that only use either topological properties or attributes.

This demonstrates the sensitivity of GNE to parameter *λ*. This parameter *λ* has a considerable impact on our method’s performance and should be appropriately selected.

#### Investigation of GNE’s predictions

We investigate the predictive ability of our model in identifying new gene interactions. For this aim, we consider two versions of BioGRID interaction datasets at two different time points (2017 and 2018 version), where the older version is used for training and the newer one is used for testing the model (temporal holdout validation). The 2018 version contains 12,835 new interactions for yeast and 11,185 new interactions for E. coli than the 2017 version. GNE’s performance trained with 50% and 80% of total interactions are comparable for both yeast and E. coli (Figure 4 and 5). We thus train our model with 50% of total interactions from the 2017 version to learn the embeddings for genes and demonstrate the impact of integrating expression data with topological properties. We create the test set with new interactions from the 2018 version of BioGRID as positive interactions and the equal number of negative interactions randomly sampled. We make predictions for these interactions using learned embeddings and create a list of (Gene *v*_*i*_, Gene *v*_*j*_, probability), ranked by the predicted probability. We consider predicted gene pairs with the probabilities of 0.5 or higher but are missing from BioGRID for further investigation as we discuss later in this section.

The temporal holdout performance of our model in comparison to other methods is shown in Figure 6. We observe that GNE outperforms both node2vec and LINE in temporal holdout validation across both yeast and E. coli datasets, indicating GNE can accurately predict new genetic interactions. Table 5 shows that GNE achieves substantial improvement of 7.0 (AUROC), 7.4 (AUPR) on yeast and 6.6 (AUROC), 5.9 (AUPR) on E. coli datasets.

**Figure 6.**
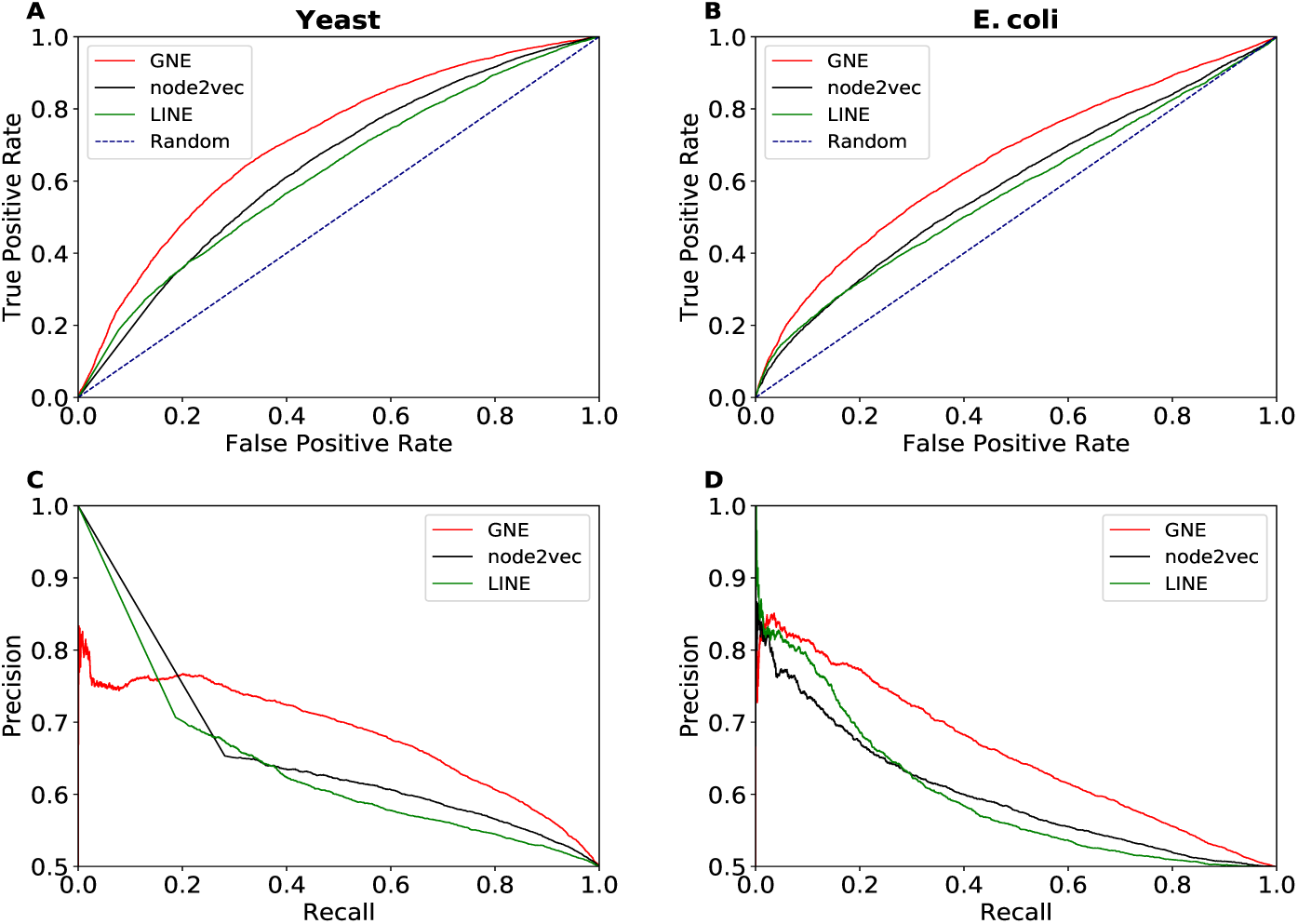
Temporal holdout validation in predicting new interactions. Performance is measured by the area under the ROC curve and the area under the precision-recall curve. Shown are the performance of each method based on the AUROC (**A, B**) and AUPR (**C, D**) for yeast and E. coli. The limit of the y-axis is adjusted to [0.5, 1.0] for the precision-recall curve to make the difference in performance more visible. GNE outperforms LINE and node2vec.

**Table 5.** AUROC and AUPR comparision for temporal holdout validation.

| Methods | Yeast |  | E. coli |  |
| --- | --- | --- | --- | --- |
|  | AUROC | AUPR | AUROC | AUPR |
| LINE | 0.620 | 0.611 | 0.569 | 0.598 |
| node2vec | 0.640 | 0.609 | 0.587 | 0.599 |
| **GNE (our model)** | **0.710** | **0.683** | **0.653** | **0.658** |

Table 6 shows the top 5 interactions with the significant increase in predicted probability for both yeast and E. coli after expression data is integrated. We also provide literature evidence with experimental evidence code obtained from the BioGRID database^26^ supporting these predictions. BioGRID compiles interaction data from numerous publications through comprehensive curation efforts. Taking new interactions added to BioGRID (version 3.4.158) into consideration, we evaluate the probability of these interactions predicted by GNE trained with and without expression data. Specifically, integration of expression data increases the probability of 8,331 (out of 11,185) interactions for E. coli (improving AUROC from 0.606 to 0.662) and 6,010 (out of 12,835) interactions for yeast (improving AUROC from 0.685 to 0.707). Integration of topology and expression data significantly increases the probabilities of true interactions between genes (Table 6).

**Table 6.** New gene interactions that are assigned high probability by GNE

| Organism | Probability |  | Gene *i* | Gene *j* | Experimental Evidence Code |
| --- | --- | --- | --- | --- | --- |
|  | Topology | Topology + Expression |  |  |  |
| Yeast | 0.287 | 0.677 | TFC8 | DHH1 | Affinity Capture-RNA<sup>34</sup> |
|  | 0.394 | 0.730 | SYH1 | DHH1 | Affinity Capture-RNA<sup>34</sup> |
|  | 0.413 | 0.746 | CPR7 | DHH1 | Affinity Capture-RNA<sup>34</sup> |
|  | 0.253 | 0.551 | MRP10 | DHH1 | Affinity Capture-RNA<sup>34</sup> |
|  | 0.542 | 0.835 | RPS13 | ULP2 | Affinity Capture-MS<sup>35</sup> |
| E. coli | 0.014 | 0.944 | ATPB | RFBC | Affinity Capture-MS<sup>36</sup> |
|  | 0.012 | 0.941 | NARQ | CYDB | Affinity Capture-MS<sup>36</sup> |
|  | 0.013 | 0.937 | PCNB | PAND | Affinity Capture-MS<sup>36</sup> |
|  | 0.015 | 0.939 | FLIF | CHEY | Affinity Capture-MS<sup>36</sup> |
|  | 0.017 | 0.938 | YCHM | PROB | Affinity Capture-MS<sup>36</sup> |

New gene interactions on 2018 version that are assigned high probability by GNE after integration of expression data. We provide probability predicted by GNE (with/without expression data) for new interactions in the 2018 version and evidence supporting the existence of predicted interactions.

To further evaluate GNE’s predictions, we consider the new version of BioGRID (version 3.4.162) and evaluate 2,609 yeast gene pairs (Additional file 1 Table S1) and 871 E. coli gene pairs (Additional file 1 Table S2) predicted by GNE with the probabilities of 0.5 or higher. We find that 128 (5%) yeast gene pairs and 78 (9%) E. coli gene pairs are true interactions that have been added to the latest release of BioGRID. We then evaluate the predictive ability of GNE by calculating the percentage of true interactions with regard to different probability bins (Figure 7). 16% of predicted yeast gene pairs and 17.5% of predicted E. coli gene pairs with the probability higher than 0.9 are true interactions. This suggests that gene pairs with high probability predicted by GNE are more likely to be true interactions.

**Figure 7.**
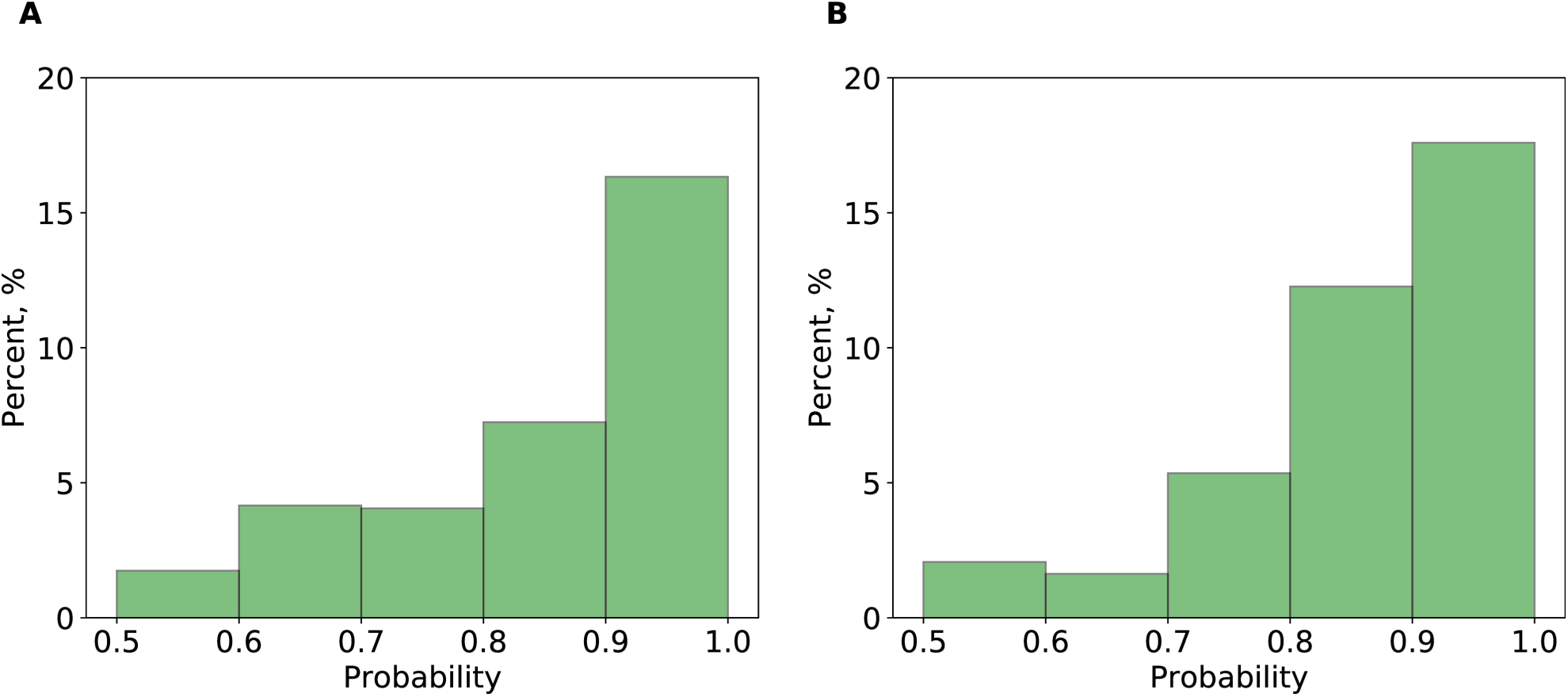
The percentage of true interactions from GNE’s predictions with different probability bins. (A) yeast (B) E. coli. We divide the gene pairs based on their predicted probabilities to different probability ranges (as shown in the x-axis) and identify the number of predicted true interactions in each range. Each bar indicates the percentage of true interactions out of predicted gene pairs in that probability range.

To support our finding that GNE predicted gene pairs have high value, we manually check gene pairs that have high predicted probability but are missing from the latest BioGRID release. We find that these gene pairs interact with the same set of other genes. For example, GNE predicts the interaction between YDR311W and YGL122C with the probability of 0.968. Mining BioGRID database, we find that these genes interact with the same set of 374 genes. Similarly, E. coli genes DAMX and FLIL with the predicted probability of 0.998 share 320 interacting genes. In this way, we identify all interacting genes shared by each of the predicted gene pairs in yeast and E. coli (Additional file 1 Table S1 and S2). Figure 8 shows the average number of interacting genes shared by a gene pair. In general, gene pairs with a high GNE probability tend to have a large number of interacting genes. For example, gene pairs with the probability greater than 0.9 have, on average, 82 common interacting genes for yeast and 58 for E. coli. Two sample t-test analysis has shown that there is a significant difference in the number of shared interacting genes with respect to different probability bins (Table 7).

**Figure 8.**
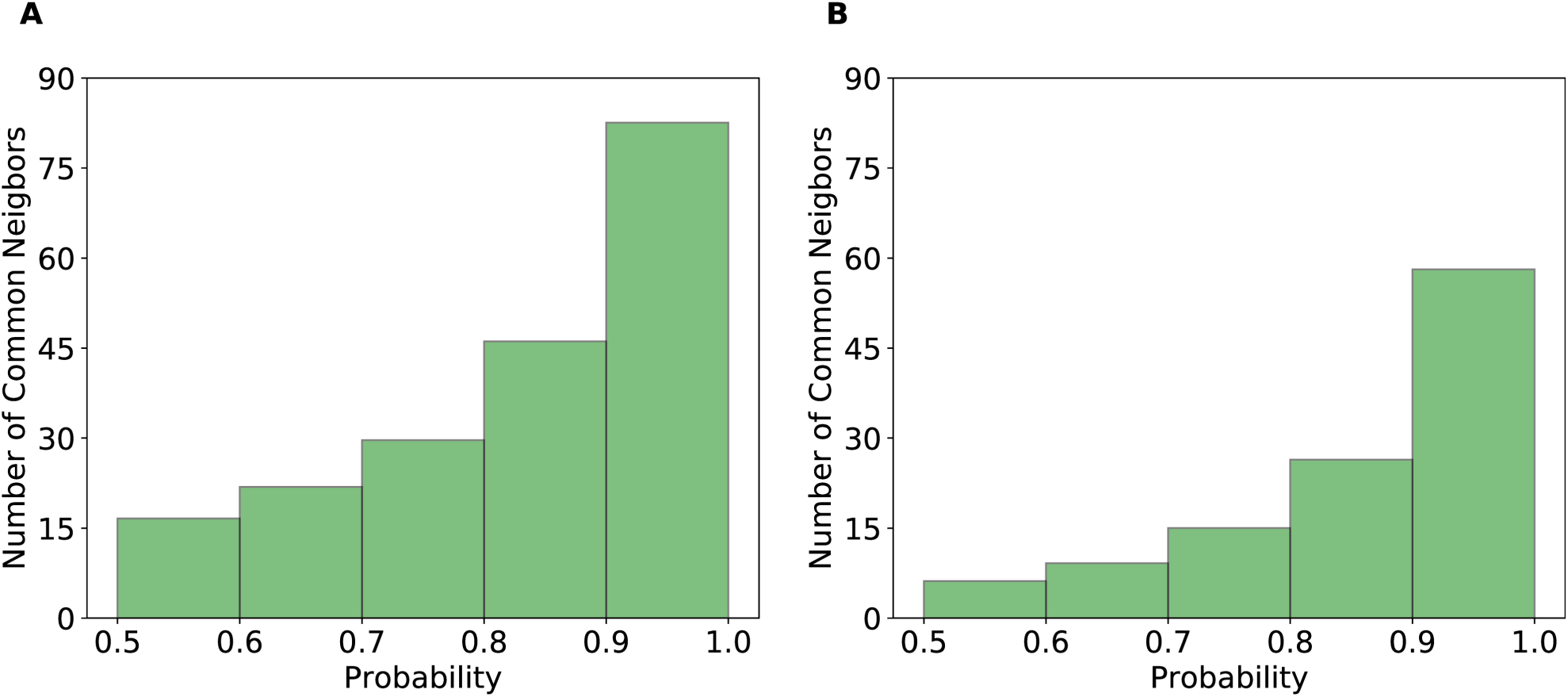
The average number of common interacting genes between the gene pairs predicted by GNE. (A) yeast (B) E. coli. We divide gene pairs into different probability groups based on predicted probabilities by GNE and compute the number of common interacting genes shared by these gene pairs. We categorize these gene pairs into different probability ranges (as shown in the x-axis). Each bar represents the average number of common interacting genes shared by gene pairs in each probability range.

**Table 7.**
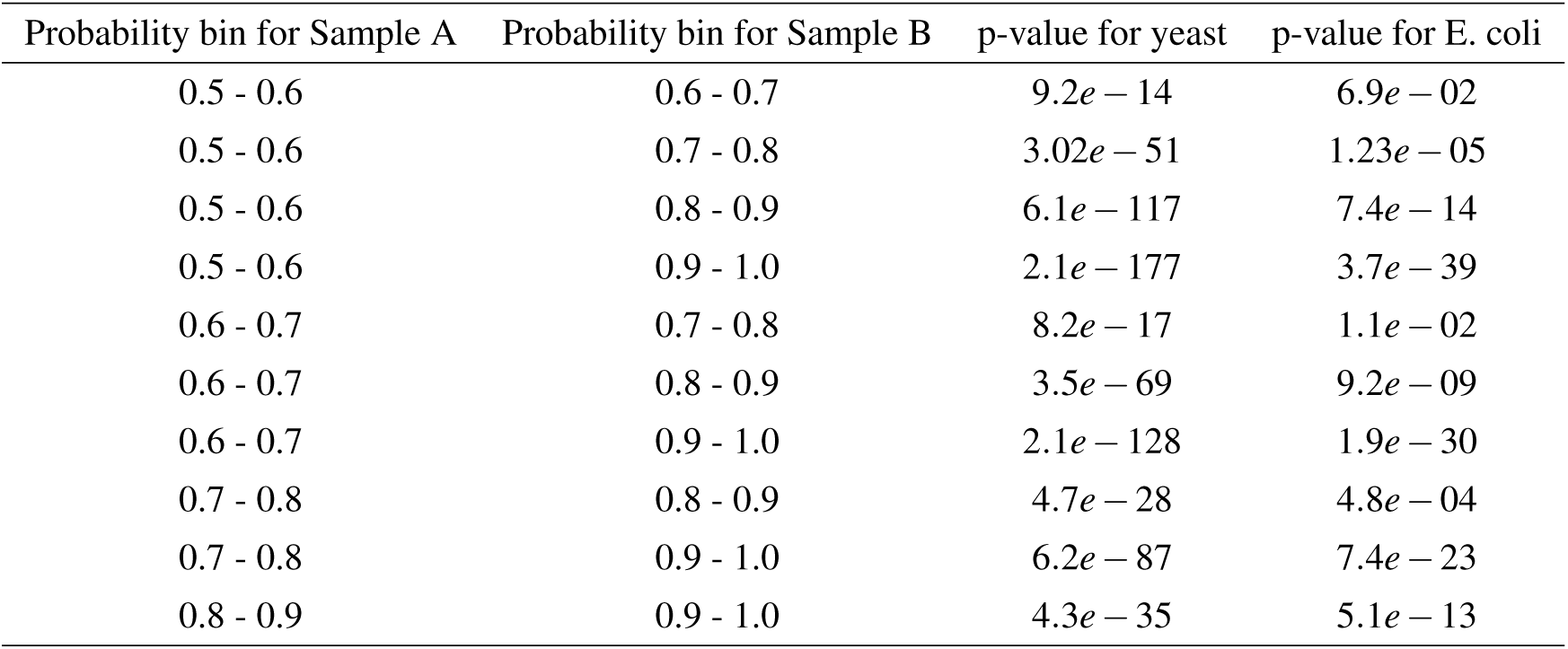
Results of two-sample t-test. We divide gene pairs into different probability groups based on predicted probabilities by GNE and compute the number of common interacting genes shared by these gene pairs. Significance test shows there is the significant difference between average number of shared genes in different probability bins.

| Probability bin for Sample A | Probability bin for Sample B | p-value for yeast | p-value for E. coli |
| --- | --- | --- | --- |
| 0.5 - 0.6 | 0.6 - 0.7 | $9.2e - 14$ | $6.9e - 02$ |
| 0.5 - 0.6 | 0.7 - 0.8 | $3.02e - 51$ | $1.23e - 05$ |
| 0.5 - 0.6 | 0.8 - 0.9 | $6.1e - 117$ | $7.4e - 14$ |
| 0.5 - 0.6 | 0.9 - 1.0 | $2.1e - 177$ | $3.7e - 39$ |
| 0.6 - 0.7 | 0.7 - 0.8 | $8.2e - 17$ | $1.1e - 02$ |
| 0.6 - 0.7 | 0.8 - 0.9 | $3.5e - 69$ | $9.2e - 09$ |
| 0.6 - 0.7 | 0.9 - 1.0 | $2.1e - 128$ | $1.9e - 30$ |
| 0.7 - 0.8 | 0.8 - 0.9 | $4.7e - 28$ | $4.8e - 04$ |
| 0.7 - 0.8 | 0.9 - 1.0 | $6.2e - 87$ | $7.4e - 23$ |
| 0.8 - 0.9 | 0.9 - 1.0 | $4.3e - 35$ | $5.1e - 13$ |

Moreover, we search the literature to see if we can find supporting evidence for predicted interactions. We find literature evidence for an interaction between YCL032W (STE50) and YDL035C (GPR1), which has the probability of 0.98 predicted by GNE. STE50 is an adaptor that links G-protein complex in cell signalling, and GPR1 is a G-protein coupled receptor. Both STE50 and GPR1 share a common function of cell signalling via G-protein. Besides, STE50p interacts with STE11p in the two-hybrid system, which is a cell-based system examining protein-protein interactions^37^. Also, BioGRID has evidence of 30 physical and 4 genetic associations between STE50 and STE11. Thus, STE50 is highly likely to interact with STE11, which in turn interacts with GPR1.

This analysis demonstrates the potential of our method in the discovery of gene interactions. Also, GNE can help the curator to identify interactions with strong potential that need to be looked at with experimental validation or within the literature.

#### Conclusion

We developed a novel deep learning framework, namely GNE to perform gene network embedding. Specifically, we design deep neural network architecture to model the complex statistical relationships between gene interaction network and expression data. GNE is flexible to the addition of different types and number of attributes. The features learned by GNE allow us to use out-of-the-box machine learning classifiers like Logistic Regression to predict gene interactions accurately.

GNE relies on a deep learning technique that can learn the underlying patterns of gene interactions by integrating heterogeneous data and extracts features that are more informative for interaction prediction. Experimental results show that GNE achieve better performance in gene interaction prediction over other baseline approaches in both yeast and E. coli organisms. Also, GNE can help the curator to identify the interactions that need to be looked at.

As future work, we aim to study the impact of integrating other sources of information about gene such as transcription factor binding sites, functional annotations (from gene ontology), gene sequences, metabolic pathways, etc. into GNE in predicting gene interaction.

## 2 Declarations

### Availability of data and material

The datasets generated and/or analysed during the current study are available.

## 3 Additional Files

### Additional file 1: Supplementary Tables

**Table S1** includes yeast gene pairs predicted by GNE with probabilities of 0.5 or higher. **Table S2** includes E. coli gene pairs predicted by GNE with probabilities of 0.5 or higher. Rows marked with yellow color indicate predicted interaction is true based on latest version 3.4.162 of BioGRID interaction dataset released on June 2018. Supplementary materials are available at http://kishankc.com.np/GNE_results/.

## References

1. Mani, R., Onge, R. P. S., Hartman, J. L., Giaever, G. & Roth, F. P. Defining genetic interaction. Proc. Natl. Acad. Sci. 105, 3461–3466 (2008).

2. Boucher, B. & Jenna, S. Genetic interaction networks: better understand to better predict. Front. genetics 4, 290 (2013).

3. Lage, K. Protein–protein interactions and genetic diseases: the interactome. Biochimica et Biophys. Acta (BBA)-Molecular Basis Dis. 1842, 1971–1980 (2014).

4. Madhukar, N. S., Elemento, O. & Pandey, G. Prediction of genetic interactions using machine learning and network properties. Front. bioengineering biotechnology 3, 172 (2015).

5. Oliver, S. Proteomics: guilt-by-association goes global. Nature 403, 601 (2000).

6. Cho, H., Berger, B. & Peng, J. Compact integration of multi-network topology for functional analysis of genes. Cell systems 3, 540–548 (2016).

7. Marbach, D. et al. Wisdom of crowds for robust gene network inference. Nat. methods 9, 796 (2012).

8. Li, R., KC, K., Cui, F. & Haake, A. R. Sparse covariance modeling in high dimensions with gaussian processes. In Proceedings of The 32nd Conference on Neural Information Processing Systems (NIPS) (2018).

9. Cui, P., Wang, X., Pei, J. & Zhu, W. A survey on network embedding. arXiv preprint arXiv:1711.08752 (2017).

10. Lei, Y.-K., You, Z.-H., Ji, Z., Zhu, L. & Huang, D.-S. Assessing and predicting protein interactions by combining manifold embedding with multiple information integration. In BMC bioinformatics, vol. 13, S3 (BioMed Central, 2012).

11. Alanis-Lobato, G., Cannistraci, C. V. & Ravasi, T. Exploitation of genetic interaction network topology for the prediction of epistatic behavior. Genomics 102, 202–208 (2013).

12. Tenenbaum, J. B., De Silva, V. & Langford, J. C. A global geometric framework for nonlinear dimensionality reduction. science 290, 2319–2323 (2000).

13. Mikolov, T., Sutskever, I., Chen, K., Corrado, G. S. & Dean, J. Distributed representations of words and phrases and their compositionality. In Advances in neural information processing systems, 3111–3119 (2013).

14. Grover, A. & Leskovec, J. node2vec: Scalable feature learning for networks. In Proceedings of the 22nd ACM SIGKDD international conference on Knowledge discovery and data mining, 855–864 (ACM, 2016).

15. Tu, Y., Stolovitzky, G. & Klein, U. Quantitative noise analysis for gene expression microarray experiments. Proc. Natl. Acad. Sci. 99, 14031–14036 (2002).

16. Tang, J. et al. Line: Large-scale information network embedding. In Proceedings of the 24th International Conference on World Wide Web, 1067–1077 (International World Wide Web Conferences Steering Committee, 2015).

17. Snoek, C. G., Worring, M. & Smeulders, A. W. Early versus late fusion in semantic video analysis. In Proceedings of the 13th annual ACM international conference on Multimedia, 399–402 (ACM, 2005).

18. He, K., Zhang, X., Ren, S. & Sun, J. Deep residual learning for image recognition. In Proceedings of the IEEE conference on computer vision and pattern recognition, 770–778 (2016).

19. Pennington, J., Socher, R. & Manning, C. Glove: Global vectors for word representation. In Proceedings of the 2014 conference on empirical methods in natural language processing (EMNLP), 1532–1543 (2014).

20. Levy, O., Goldberg, Y. & Dagan, I. Improving distributional similarity with lessons learned from word embeddings. Transactions Assoc. for Comput. Linguist. 3, 211–225 (2015).

21. Kingma, D. P. & Ba, J. Adam: A method for stochastic optimization. arXiv preprint arXiv:1412.6980 (2014).

22. Duchi, J., Hazan, E. & Singer, Y. Adaptive subgradient methods for online learning and stochastic optimization. J. Mach. Learn. Res. 12, 2121–2159 (2011).

23. Tieleman, T. & Hinton, G. Lecture 6.5-rmsprop, coursera: Neural networks for machine learning. Univ. Toronto, Tech. Rep. (2012).

24. Srivastava, N., Hinton, G., Krizhevsky, A., Sutskever, I. & Salakhutdinov, R. Dropout: A simple way to prevent neural networks from overfitting. The J. Mach. Learn. Res. 15, 1929–1958 (2014).

25. Ioffe, S. & Szegedy, C. Batch normalization: Accelerating deep network training by reducing internal covariate shift. arXiv preprint arXiv:1502.03167 (2015).

26. Stark, C. et al. Biogrid: a general repository for interaction datasets. Nucleic acids research 34, D535–D539 (2006).

27. Butte, A. & Hohane, L. Mutual information relevance networks: functional genomic clustering using pairwise entropy measurements. (2000).

28. Chua, H. N., Sung, W.-K. & Wong, L. Exploiting indirect neighbours and topological weight to predict protein function from protein–protein interactions. Bioinformatics 22, 1623–1630 (2006).

29. Abadi, M. et al. TensorFlow: Large-scale machine learning on heterogeneous systems (2015). Software available from tensorflow.org.

30. Clevert, D.-A., Unterthiner, T. & Hochreiter, S. Fast and accurate deep network learning by exponential linear units (elus). arXiv preprint arXiv:1511.07289 (2015).

31. Davis, J. & Goadrich, M. The relationship between precision-recall and roc curves. In Proceedings of the 23rd international conference on Machine learning, 233–240 (ACM, 2006).

32. Maaten, L. v. d. & Hinton, G. Visualizing data using t-sne. J. machine learning research 9, 2579–2605 (2008).

33. Mao, F., Dam, P., Chou, J., Olman, V. & Xu, Y. Door: a database for prokaryotic operons. Nucleic acids research 37, D459–D463 (2008).

34. Miller, J. E. et al. Genome-wide mapping of decay factor–mrna interactions in yeast identifies nutrient-responsive transcripts as targets of the deadenylase ccr4. G3: Genes, Genomes, Genet. 8, 315–330 (2018).

35. Liang, J. et al. Recruitment of a sumo isopeptidase to rdna stabilizes silencing complexes by opposing sumo targeted ubiquitin ligase activity. Genes & development 31, 802–815 (2017).

36. Babu, M. et al. Global landscape of cell envelope protein complexes in escherichia coli. Nat. biotechnology 36, 103 (2018).

37. Gustin, M. C., Albertyn, J., Alexander, M. & Davenport, K. Map kinase pathways in the yeastsaccharomyces cerevisiae. Microbiol. Mol. biology reviews 62, 1264–1300 (1998).

